# Mitochondrial glycerol phosphate oxidation is modulated by adenylates through allosteric regulation of cytochrome *c* oxidase activity in mosquito flight muscle

**DOI:** 10.1101/624452

**Authors:** Alessandro Gaviraghi, Juliana B.R. Correa Soares, Julio A. Mignaco, Carlos Frederico L. Fontes, Marcus F. Oliveira

**Affiliations:** Laboratório de Bioquímica de Resposta ao Estresse, Instituto de Bioquímica Médica Leopoldo de Meis, Universidade Federal do Rio de Janeiro, Cidade Universitária, Rio de Janeiro, RJ, Brazil; Instituto Nacional de Ciência e Tecnologia em Entomologia Molecular (INCT-EM), Rio de Janeiro, RJ, Brazil; Laboratório de Estrutura e Regulação de Proteínas e ATPases, Instituto de Bioquímica Médica Leopoldo de Meis, Universidade Federal do Rio de Janeiro, Cidade Universitária, Rio de Janeiro, RJ, Brazil

**Keywords:** metabolism, bioenergetics, respiration, oxidative phosphorylation, electron transport chain

## Abstract

The huge energy demand posed by insect flight activity is met by an efficient oxidative phosphorylation process that takes place within flight muscle mitochondria. In the major arbovirus vector *Aedes aegypti*, mitochondrial oxidation of pyruvate, proline and glycerol 3 phosphate (G3P) represent the major energy sources of ATP to sustain flight muscle energy demand. Although adenylates exert critical regulatory effects on several mitochondrial enzyme activities, the potential consequences of altered adenylate levels to G3P oxidation remains to be determined. Here, we report that mitochondrial G3P oxidation is controlled by adenylates through allosteric regulation of cytochrome c oxidase (COX) activity in *A. aegypti* flight muscle. We observed that ADP significantly activated respiratory rates linked to G3P oxidation, in a protonmotive force-independent manner. Kinetic analyses revealed that ADP activates respiration through a slightly cooperative mechanism. Despite adenylates caused no effects on G3P-cytochrome *c* oxidoreductase activity, COX activity was allosterically activated by ADP. Conversely, ATP exerted powerful inhibitory effects on respiratory rates linked to G3P oxidation and on COX activity. We also observed that high energy phosphate recycling mechanisms did not contribute to the regulatory effects of adenylates on COX activity or G3P oxidation. We conclude that mitochondrial G3P oxidation by *A. aegypti* flight muscle is regulated by adenylates essentially through the allosteric modulation of COX activity, underscoring the bioenergetic relevance of this novel mechanism and the potential consequences for mosquito dispersal.

## Introduction

Adult females of *Aedes aegypti* are hematophagous insects and vectors of several arboviruses to humans, including Dengue, Zika, Chikungunya and yellow fever viruses. The explosive spreading of arboviral infections observed in tropical and subtropical countries, represents a major public health problem, affecting nearly 400 million individuals each year with 3.9 billion people at risk in 128 countries [1–3]. Given the absence of vaccines for most of these viruses, vector population control remains the main strategy for preventing or reducing arbovirus transmission. Dispersal capacity of insect vectors through flight activity represents an important parameter for vector-borne disease control strategies. The contractile activity of flight muscles located in vector thoraces provides the workload to sustain flight activity, which poses a huge energy demand to these organisms. Indeed, insect flight muscles were reportedly found to be one of the most metabolically active tissues found in nature [4], exhibiting up to 100-fold increase in respiratory rates in the transition between rest and flight [5–8]. The energy required to sustain flight among different insect species is essentially provided by a very efficient oxidative phosphorylation process that takes place within mitochondria. Observations from different laboratories have demonstrated the unique structural and functional properties of flight muscle mitochondria [5,9–11]. A remarkable feature is the extreme packaging of mitochondrial inner membrane in insect flight muscle, which is consistent with the high respiratory capacity and ATP production rates [10–14].

A key property of aerobic organisms is the control of the rates of oxidative phosphorylation according to the cell energy demand. This represents the basis of the so-called respiratory control, which can be a consequence of: *i)* regulation of the *pmf* by ADP availability [15,16], which allows ATP synthesis through chemiosmosis [17], or *ii)* regulation of mitochondrial enzymes including isocitrate dehydrogenase [18–20], proline dehydrogenase [21], and cytochrome *c* oxidase (COX) activity by ADP/ATP levels [22,23]. The first control mechanism is abolished by the use of proton ionophores that dissipate *pmf*, which increases respiratory rates independent of the energy charge. On the other hand, the second mechanism occurs independently of *pmf*, and is based on the ability of adenylates to allosterically modulate mitochondrial dehydrogenases and COX activities by direct binding of ATP or ADP [22,23]. Regulation of COX activity by adenylates is particularly relevant to bioenergetics given its key role in electron transport system to allow respiration and ATP synthesis.

COX is the terminal enzyme complex of the electron transport system and catalyzes the transfer of four electrons from cytochrome *c* (cyt*c*) to molecular oxygen, which is reduced to two molecules of water. This transmembrane complex is located at the inner mitochondrial membrane and, in mammals, is composed by 13 different subunits [24,25]. The catalytic core of this complex is composed by three subunits encoded by mitochondrial-DNA, while the remaining ten subunits encoded by nuclear-DNA are located around the core and play a role in stabilizing and regulating COX function [26,27]. Evidence that adenylates would regulate electron flow at COX demonstrate that at physiological concentrations, ATP would bind to this complex, decreasing its affinity to cyt*c* [28]. Further evidence demonstrated the existence of several ATP and ADP binding sites in different COX subunits [29–31]. Indeed, COX from bovine heart binds with high-affinity ten ADP molecules, seven of which are exchanged by ATP at high ATP/ADP ratios [32]. Particularly relevant are subunits IV and VIa, which have been identified as the most important adenylate binding sites, with a regulatory effect on COX activity [22,23,29,32–35].

Despite the medical importance of *Aedes aegypti*, few studies investigated the biological relationships between mitochondrial physiology and mosquito biology. Previous evidence from our laboratory revealed that blood feeding caused transient reductions in oxygen consumption and hydrogen peroxide generation in *A. aegypti* flight muscle mitochondria [10]. These changes were parallel to altered mitochondrial dynamics, towards organelle fusion upon blood feeding. Further investigations revealed that respiration was essentially sustained by the oxidation of pyruvate, proline, and glycerol 3 phosphate (G3P) [11], which promoted distinct bioenergetic capacities, but with preserved efficiencies [11], as schematically outlined in figure 1.

**Figure 1.**
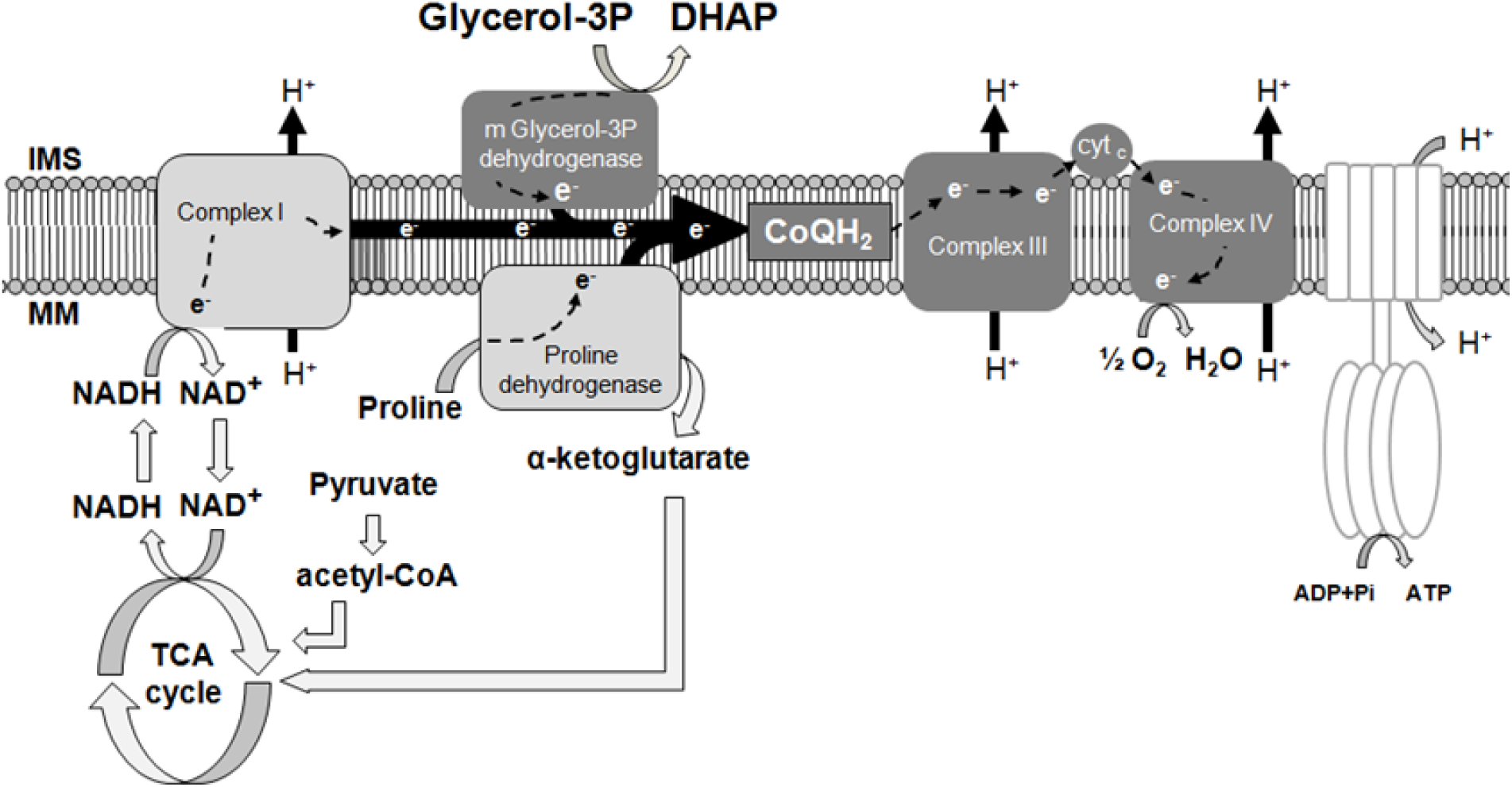
Schematic representation of the electron transport system in *A. aegypti* mitochondria and the main substrates that contribute to oxidative phosphorylation. The two main sites of electron entry into the system are mediated by pyruvate and proline (through complex I and proline dehydrogenase path, light grey boxes), and through glycerol 3 phosphate (G3P, through mitochondrial glycerol 3 phosphate dehydrogenase path, dark grey boxes), as previously described [11]. Reduced ubiquinone (CoQH_2_), complex III, cytochrome *c* (cyt*c*), and complex IV represent a convergent common path for both substrates. IMS, intermembrane space, MM, mitochondrial matrix, DHAP, dihydroxyacetone phosphate.

G3P can be synthesized by either phosphorylation of glycerol released from triacylglycerol hydrolysis by glycerol kinase, or by reduction of dihydroxyacetone phosphate (DHAP) derived from glycolysis by the cytosolic isoform of G3P dehydrogenase (cG3PDH). The mitochondrial isoform of G3P dehydrogenase (mG3PDH) oxidizes G3P back to DHAP and then transfers the cytosolic reducing equivalents directly to the electron transport system [36]. Therefore, the concerted action of cG3PDH and mG3PDH creates a mechanism that connects glycolysis/triacylglycerol metabolism to mitochondrial electron transport system through the glycerophosphate shuttle. Interestingly, the glycerophosphate shuttle activity is remarkably high in tissues with high energy demand such as in mammalian brown adipose tissue, [37–39]. Also, the intense G3P production relative to lactate during insect flight indicate that glycerophosphate shuttle has a prominent role to recycle cytosolic NAD+ in insects to sustain intense glycolytic flux than lactate dehydrogenase [40,41]. Although mG3PDH is strongly regulated by hormones, calcium [42–44], and inhibited by metformin [45], the potential effect of adenylates on mitochondrial G3P oxidation remains to be determined. To fill this gap of knowledge, we postulated that respiratory rates sustained by G3P oxidation in *A. aegypti* would be strongly controlled by adenylates at the level of COX activity. This regulatory mechanism would be particularly relevant not only to allow prompt energy provision for myofibril contraction when flight is required, but also to avoid unnecessary substrate consumption during resting periods. Here, we demonstrate that *A. aegypti* flight muscle respiratory rates sustained by G3P oxidation were stimulated by ADP, acting as an allosteric modulator, with moderate cooperativity, independently of substrate transport to mitochondria, tricarboxylic acid cycle enzyme activities, and *pmf*. Modulating effects of adenylates were specific to COX activity and not to other ETS components involved in G3P oxidation. Conversely, ATP exerted powerful inhibitory effects on mitochondrial G3P oxidation and COX activity. We conclude that respiration sustained by G3P oxidation in *A. aegypti* flight muscle mitochondria is strongly regulated by adenylates essentially through the allosteric modulation of COX activity, involving a direct bioenergetic crosstalk between mitochondria and myofibrils, and with potential consequences to mosquito flight and dispersal.

## Results

### Adenylates regulate respiratory rates independent of *pmf*, but with magnitudes dependent on fuel preference

The general architecture of female *A. aegypti* flight muscle revealed that myofibrils were intercalated with threads of giant mitochondria [10], with highly packed and abundant cristae as previously reported for other insect models [46]. The observation of these remarkable architectural features in flight muscle suggest that substrates exchange might be favored between the energy supply (mitochondria) and demand (myofibrils) sites to sustain intense flight mechanical work. In this sense, our laboratory demonstrated that respiratory rates in *A. aegypti* flight muscle are essentially sustained by the oxidation of pyruvate, proline, and G3P [11], as schematically outlined in figure 1. Evidence demonstrate that multiple signals can regulate mitochondrial G3P dehydrogenase (mG3PDH), with direct impacts on respiratory rates [42–44]. However, how respiratory rates sustained by G3P oxidation in mitochondria are regulated by adenylates remains to be determined. To address this, we firstly determined oxygen consumption rates in mechanically permeabilized flight muscle and isolated mitochondria from female insects in the presence or the absence of 2 mM ADP using pyruvate plus proline (Pyr+Pro) or G3P as substrates.

Maximum uncoupled respiratory rates in both mechanically permeabilized flight muscle and isolated mitochondria were significantly higher in both substrates when 2 mM ADP was present (Figure 2 and S1). Important differences emerged when we compared the effects of ADP on respiratory rates sustained by Pyr+Pro (Figures 2A and S1A) with those produced by G3P oxidation (Figures 2B and S1B). While 2 mM ADP boosted respiratory rates sustained by Pyr+Pro from 6.3 to 11 times (Figures 2A, and S1A), the stimulatory effect of ADP under G3P oxidation was in the range of 50 - 120% (Figures 2B and S1B). The differential responses of substrate oxidation to ADP can be ascribed to the known regulatory effects of ADP on multiple mitochondrial dehydrogenases [18,21,47,48], which exert a higher effect on complex I-dependent substrates including pyruvate and proline, and also on COX [22,23]. We also observed that 2 mM ATP in the presence of an ATP regenerating system caused a strong inhibition (∼62%) of respiratory rates provided by G3P in permeabilized flight muscle (Figure 2B). Given that mitochondrial G3P oxidation occurs independently of mitochondria inner membrane transport and TCA cycle dehydrogenases, and no direct effect of adenylates on G3PDH activity was so far reported, this implicates an alternative mechanism(s) by which adenylates would regulate respiration. We then performed the subsequent experiments aiming to understand the mechanistic basis of the regulatory effects of adenylates on mitochondrial G3P oxidation in mosquito flight muscle.

**Figure 2.**
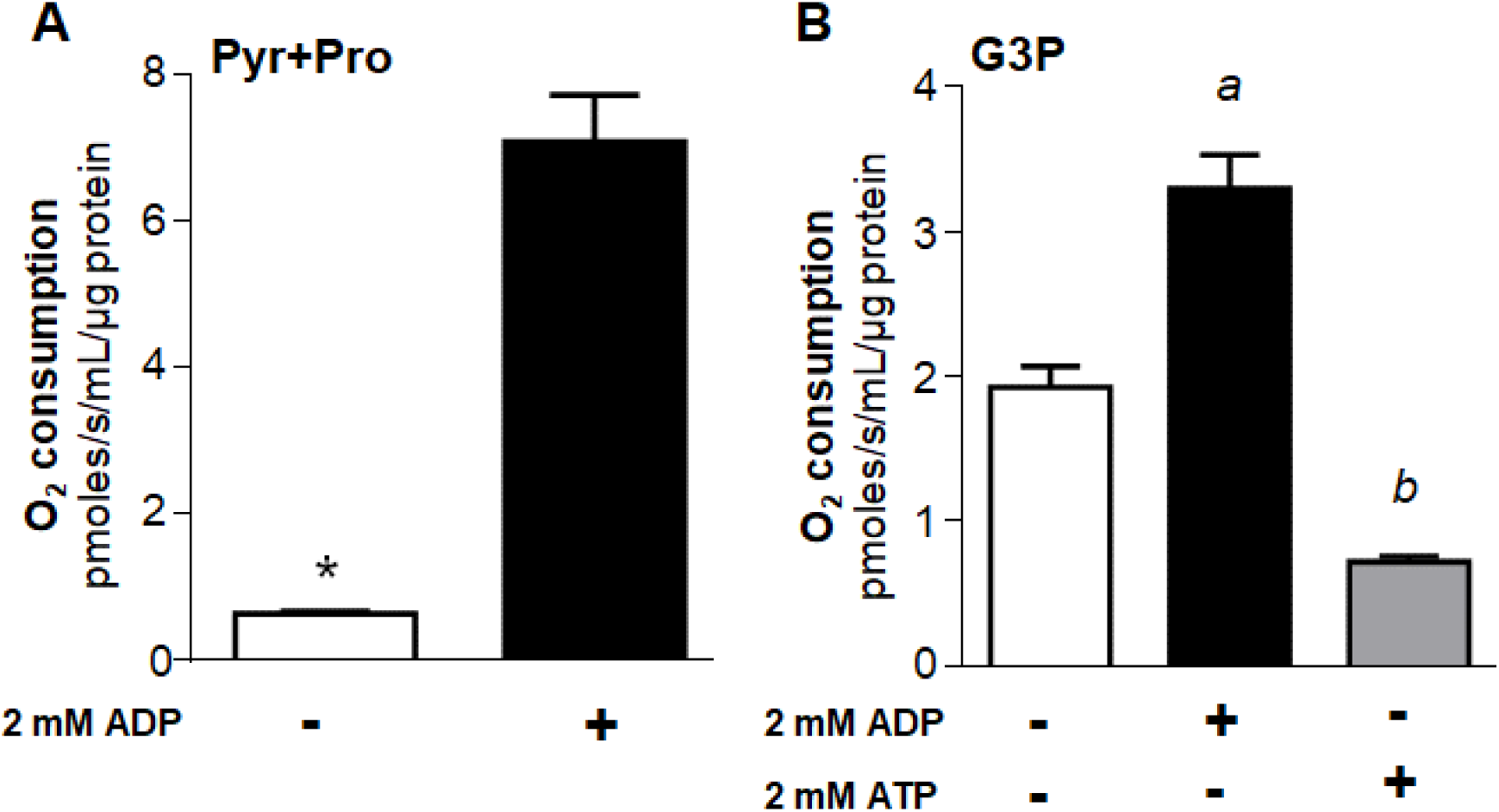
Adenylates regulate respiratory rates by a mechanism independent of mitochondrial substrate transport, tricarboxylic acid cycle enzymes activities, and protonmotive force. Maximum uncoupled respiratory rates (2.5 µM FCCP) of mechanically permeabilized flight muscle from adult females were determined by high resolution respirometry in the absence (−) or in the presence (+) of 2 mM ADP using 10 mM pyruvate plus proline (A), or 20 mM G3P (B) as substrates. When using G3P, 0.5 µM rotenone was added before G3P to avoid electron backflow from G3PDH to complex I. The effect of 2 mM ATP + regenerating system was also assessed in permeabilized flight muscle using G3P as a substrate (B). Data are expressed as mean O_2_ consumption rates (pmoles/s/mL/µg protein) ± standard error of the mean (SEM) of at least four different experiments. Comparisons between groups were done by unpaired Student’s t test with \**p*<0.0001 relative to control group (without ADP), or by ANOVA and *a posteriori* Tukey’s tests with ^a^ *p*<0.0001 relative to control and ATP, ^b^ *p*<0.005 relative to control.

### ADP activates mitochondrial G3P oxidation in either isolated organelles or permeabilized flight muscle through a slight cooperative mechanism

Our next step was to compare the effects of ADP titration on G3P-linked respiratory rates using mechanically permeabilized flight muscle and isolated mitochondria (Figure 3 and Table 1).

**Figure 3.**
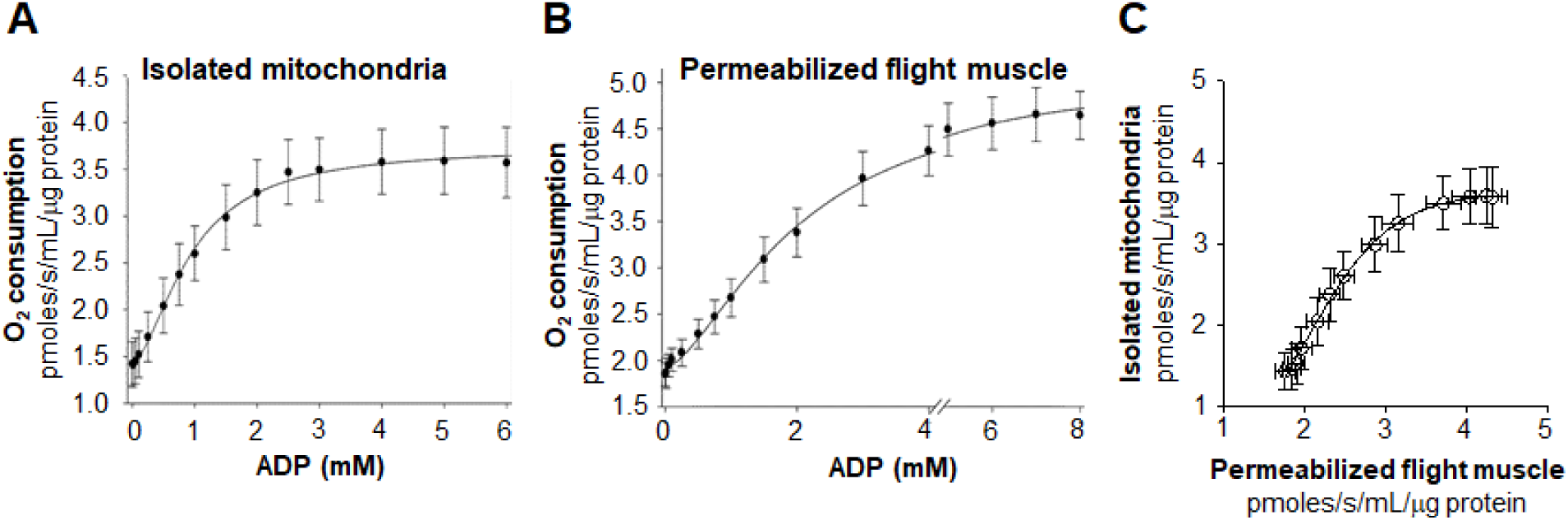
Activation of respiratory rates by ADP in isolated mitochondria and mechanically permeabilized flight muscle. Saturation kinetic curves were determined under uncoupling conditions (2.5 µM FCCP) on isolated mitochondria (200 µg protein, A) or mechanically permeabilized flight muscle (single flight muscle, B) by high resolution respirometry using ADP concentrations from 0 to up to 8 mM in the presence of 0.5 µM rotenone + 20 mM G3P. Data were fitted using the Hill four parameters equation, and are expressed as mean O_2_ consumption rates (pmoles/s/mL/µg protein) ± standard error of the mean (SEM) of at least four different experiments.

**Table 1.** Kinetic assessment of mitochondrial G3P oxidation from saturation curves using ADP concentrations from 0 to up to 8 mM in uncoupled isolated mitochondria and permeabilized flight muscle. Apparent kinetic parameters including ^app^V_max_, ^app^K_0.5_, Hill coefficient (^app^n_H_), values were determined by non-linear regressions from titration curves shown in Figure 3. ^app^V_max_ values are expressed as pmoles/s/mL/µg protein, while ^app^K_0.5_ and ^app^EC_50_ as mM. Data are expressed as mean ± standard error of the mean (SEM) of at least four different experiments.

| Sample | $^{app}V_{max}$ | $^{app}K_{0.5}$ | Hill ($^{app}n_H$) |
| --- | --- | --- | --- |
| Permeabilized flight muscle | 5.05 ± 0.13 | 2.07 ± 0.12 | 1.62 ± 0.09 |
| Isolated mitochondria | 3.73 ± 0.09 | 0.94 ± 0.04 | 1.71 ± 0.12 |

We observed that ADP significantly increased uncoupled respiratory rates in isolated *A. aegypti* flight muscle mitochondria by ∼ 2.6 fold at the highest concentration (Figure 3A). Uncoupled respiratory rates were also significantly increased by ADP in permeabilized flight muscle (Figure 3B). Indeed, respiratory rates of isolated mitochondria and permeabilized flight muscle upon ADP titration were strongly correlated (Pearson R = 0.9550, *p*<0.0001) in a non-linear fashion (Figure 3C). In average, respiratory rates induced by different ADP concentrations in isolated mitochondria represent close to 90% of the rates obtained in permeabilized flight muscle. In both cases, the ADP titration curves were mildly sigmoidal in shape, and best fit analyzes indicated that sigmoidal Hill equations produced the highest goodness of fit (R^2^) values (0.9978 for isolated mitochondria, and 0.9982 for permeabilized flight muscle). Indeed, kinetic analyses shown in Table 1 demonstrate that respiratory rates in permeabilized flight muscle exhibited a slightly lower apparent affinity (^app^K_0.5_) for ADP compared to isolated mitochondria (^app^K_0.5_ = 2.07 vs. 0.94 mM ADP). On the other hand, apparent maximal uncoupled respiratory rates (^app^V_max_) were higher in permeabilized flight muscle than in isolated mitochondria (^app^V_max_= 5.05 vs. 3.73 pmoles/s/mL/µg protein) (Table 1).

These results show that the effect of ADP on G3P-linked respiration is an intrinsic property of *A. aegypti* flight muscle mitochondria and not an artifact generated by the loss of their native morphology and their interaction with other organelles during the isolation process. Therefore, given the technical and biological advantages to assess mitochondrial function in permeabilized muscle [49,50], and their quite similar responses of ADP on respiratory rates compared to isolated mitochondria, all subsequent experiments used exclusively permeabilized flight muscle samples.

### Adenylates strongly regulate mitochondrial G3P oxidation

To further investigate the mechanisms by which ADP modulates G3P-linked respiration in *A. aegypti* flight muscle, experiments of ADP titration on oxygen consumption rates coupled to kinetic analyses were performed. In figure 4A we show that ADP significantly increased respiratory rates in *A. aegypti* flight muscle by ∼ 2.3 fold at the highest concentration, regardless the coupling state. Indeed, the activating effects of ADP on maximal respiratory rates, and the apparent affinity, were remarkably higher in the uncoupled state (^app^V_max_= 5.05 pmoles/s/mL/µg protein, ^app^K_0.5_ = 2.07 mM ADP), than in the coupled state (^app^V_max_ = 3.45 pmoles/s/mL/µg protein, ^app^K_0.5_ = 3.22 mM) (Table 2). All ADP titration curves revealed a slightly sigmoidal shape and best fit analyzes indicated that Hill equations produced the highest goodness of fit (R^2^) values (0.9940 for coupled, 0.9982 for uncoupled, and 0.9970 for uncoupled + CAT). Indeed, kinetic analyses of ADP titration curves produced apparent Hill coefficients (^app^_*n*_H) ∼ 1.71 regardless the mitochondrial coupling state, suggesting an apparent positive cooperative effect of ADP on G3P-linked respiration in *A. aegypti* flight muscle. (Table 2). Altogether, we conclude that ADP activates *A. aegypti* flight muscle respiration through a slightly positive cooperative mechanism, independent of mitochondrial inner membrane transport, matrix dehydrogenases, and *pmf*.

**Table 2.**
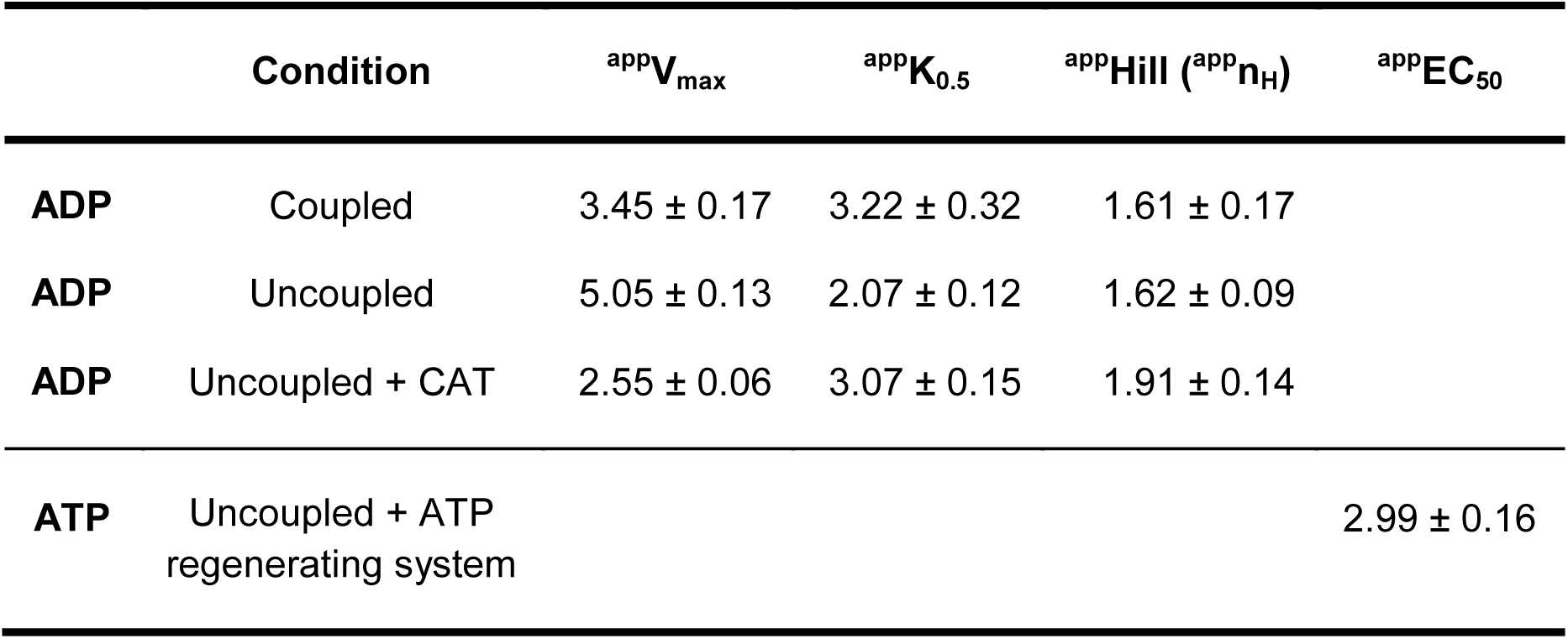
Kinetic assessment of mitochondrial G3P oxidation from saturation curves using 0 up to 10 mM ADP in coupled, uncoupled, and uncoupled + CAT conditions. The inhibitory effects of ATP on respiratory parameters were determined using 0-10 mM ATP + an ATP regenerating system in uncoupled conditions. Apparent kinetic parameters including ^app^V_max_, ^app^K_0.5_, Hill coefficient (^app^n_H_), and ^app^EC_50_ values were determined by non-linear regression from titration curves shown in Figure 4. ^app^V_max_ values are expressed as pmoles/s/mL/µg protein while ^app^K_0.5_ and ^app^EC_50_ as mM. Data are expressed as mean ± standard error of the mean (SEM) of at least four different experiments.

**Figure 4.**
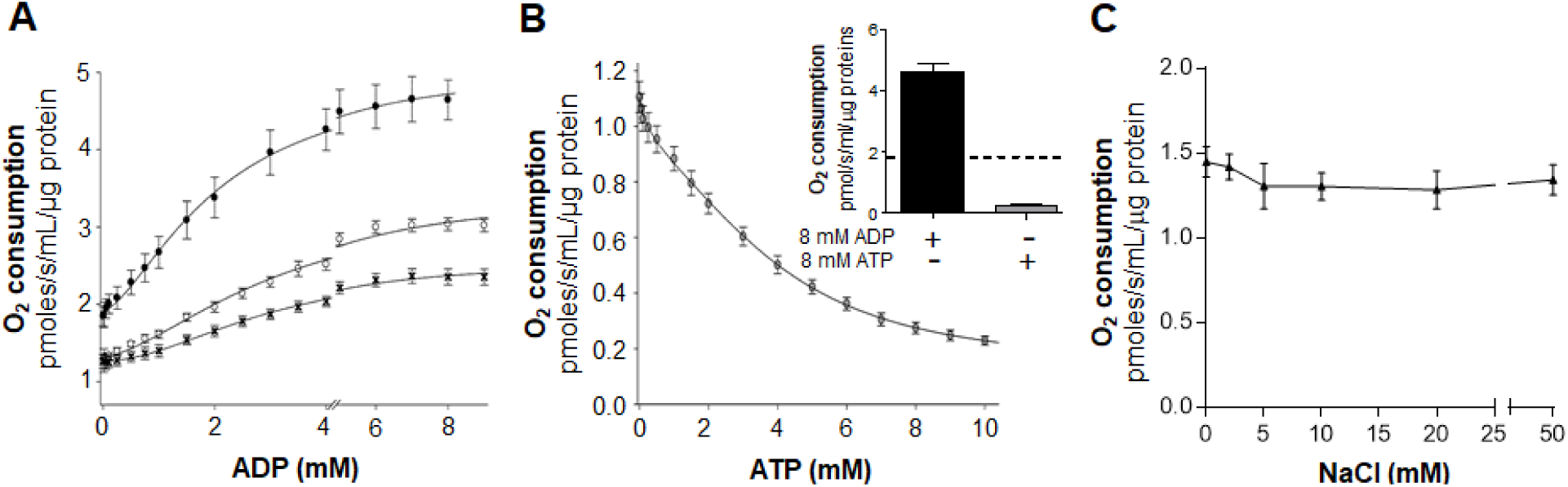
Regulatory effects of adenylates on mitochondrial G3P oxidation in *A. aegypti* flight muscle. (A) Saturation kinetics curves were determined by high resolution respirometry of mechanically permeabilized flight muscle using ADP concentrations from 0 to 9 mM in the presence of 0.5 µM rotenone + 20 mM G3P in coupled (open circles), and uncoupled (2.5 µM FCCP, black circles) conditions. Inhibition of ANT on the stimulatory effects of ADP on respiratory rates was assessed by using 50 µM CAT under uncoupled conditions (black crosses). (B) Saturation kinetics curves were also determined using ATP concentrations from 0 to 10 mM in the presence of 0.5 µM rotenone + 20 mM G3P in uncoupled (2.5 µM FCCP, grey circles) conditions in the presence of an ATP-regenerating system, as described in the methods section. The inset shows a comparison of the regulatory effects of 8 mM ADP (black bar) and 8 mM ATP (grey bar) on respiratory rates, relative to control measurements made in absence of adenylates (dashed line). Data were fitted using the Hill equation with four parameters (A) or five parameters logistic two slopes equation (B). (C) The effect of ionic strength on respiration was determined using NaCl concentrations from 0 to 50 mM in the presence of 0.5 µM rotenone + 20 mM G3P in uncoupled (2.5 µM FCCP) conditions. Comparisons between groups were done by one-way ANOVA and a posteriori Tukey’s multiple comparisons test. Data are expressed as mean O_2_ consumption rates (pmoles/s/mL/µg protein) ± standard error of the mean (SEM) of at least four different experiments.

In order to further investigate in which mitochondrial compartment ADP would regulate respiratory rates, ADP titration curves were performed under uncoupling conditions in the presence of CAT, a highly selective and potent inhibitor of the adenine nucleotide translocator (ANT) [51]. In fact, blockage of ADP-ATP exchange by the ANT reduced not only the stimulatory effects of ADP respiratory rates by ∼ 49% (^app^V_max_, 5.05 *vs.* 2.55 pmoles/s/mL/µg protein), but also the affinity for ADP (^app^K_0.5_ = 2.07 *vs.* 3.07 mM ADP) (Figure 4A, and Table 2). Despite the lower stimulatory effect of ADP on absolute respiratory rates under CAT treatment, a relative increase of ∼ 1.9 fold was observed at high *vs.* low ADP concentrations, resulting in a typical sigmoidal curve with ^app^n_H_ = 1.91 (Table 2). These data indicate that activation of respiration by ADP involves an unidentified intra and extramitochondrial regulatory components of the electron transport system in flight muscle.

To assess the effects of ATP on respiration, ATP titration curves were carried out in the presence of FCCP and an ATP-regenerating system composed of pyruvate kinase + lactate dehydrogenase. Figure 4B shows that ATP exerted a powerful inhibitory effect on G3P-linked respiration (∼79%). Best fit analyses indicated that five parameters logistic two slopes model produced the highest goodness of fit values (R^2^=0.999) with ^app^EC_50_ values ∼ 2.99 mM ATP (Table 2). Indeed, even the conversion of endogenous ADP in the presence of the ATP regenerating system, but in the absence of supplemented ATP, significantly reduced respiratory rates in the flight muscle by ∼ 40%.

Since the ADP and ATP used in our experiments were sodium salts, one might argue that alterations in ionic strength caused by higher sodium concentrations would potentially affects G3P-linked respiration. In order to rule out this possibility, uncoupled G3P-linked respiratory rates were performed in the presence of 0 to 50 mM NaCl. Figure 4C shows that increases in sodium concentrations up to 50 mM caused no apparent effects on respiration, ruling out altered ionic strength as a potential explanation for the regulatory effects of adenylates on mitochondrial G3P oxidation.

### Adenylates specifically and allosterically regulate COX activity

Since the regulatory effects of adenylates on flight muscle G3P oxidation occurs independent of mitochondrial transport, TCA cycle dehydrogenases, and *pmf*, our next step was to determine which components of the electron transport system would be targeted by ADP and ATP. To accomplish this, we determined the effects of adenylates on two distinct branches of *A. aegypti* flight muscle electron transport system: the mG3PDH–cytochrome *c* (mG3PDH-cyt*c*) oxidoreductase, and the cytochrome *c* oxidase (COX) activities. The mG3PDH-cyt*c* oxidoreductase activity was determined by following at 550 nm the rate of cyt*c* reduction induced by G3P sensitive to antimycin A. Figure 5A shows that 30 minutes pre-incubation of *A. aegypti* flight muscle homogenates with 3 mM ADP or 3 mM ATP + regenerating system caused any effect on the rate of cyt*c* reduction provided by G3P, strongly suggesting that none of the electron transport system components from mG3PDH to cyt*c* are regulated by adenylates. We then hypothesized that COX activity would be the most likely candidate for the regulatory effect of adenylates on *A. aegypti* flight muscle electron transport system. For this aim, oxygen consumption rates provided by TMPD/ascorbate were determined in permeabilized flight muscle under different concentrations of cyt*c*. Figure 5B shows that in the absence of ADP (control), COX activity fitted in a hyperbolic Michaelis-Menten-type curve, which allowed us to determine its kinetic parameters (Table 3). In the presence of 3 mM ADP, the resulting curves were also hyperbolic in shape and fitted in the Michaelis-Menten equation. Kinetic analyses revealed that ADP increased ^app^V_max_ of COX activity by ∼42% compared to control conditions (64.19 *vs.* 45.07 pmoles/s/mL/µg protein), while the apparent affinity (^app^K_m_) were quite similar between the groups (40.7 *vs.* 35.5 µM cyt*c*, Table 3). Conversely, 3 mM ATP + ATP-regenerating system caused a strong (∼74.9%) inhibitory effect on COX activity compared to control (^app^V_max_ 11.3 *vs.* 45.07 pmoles/s/mL/µg protein). Remarkably, the presence of ATP shifted the kinetics of COX activity towards a typical sigmoidal-shaped curve (Figure 5B), indicating an apparent cooperativity between the cyt*c* binding sites (^app^n_H_ ∼ 2.2, Table 3). The effects of adenylates on COX activity in flight muscle homogenate samples were also determined by an independent method (spectrophotometric quantification of cyanide-sensitive cyt*c* oxidation). Figure 5C shows that 3 mM ADP increased COX activity by ∼ 29%, while 3 mM ATP + ATP-regenerating system significantly inhibited COX activity by ∼70% compared to control. To exclude that the effect on COX activity was influenced by ionic strength, oxygen consumption rates provided by TMPD/ascorbate were determined in permeabilized flight muscle under different concentrations of NaCl. Figure 5D shows that ionic strength caused no effect on COX activity up to 50 mM NaCl. These results strengthen our observation that adenylates exert strong regulatory effects on COX activity in *A. aegypti* flight muscle.

**Table 3.** Kinetic assessment of COX activity from cyt*c* saturation curves in permeabilized flight muscle. The effects of adenylate on COX kinetic parameters were obtained from the saturation curves shown in Figure 5B. Apparent kinetic parameters including ^app^V_max_, ^app^K_m_, ^app^K_0.5_, and Hill coefficient (^app^n_H_) were determined graphically from titration curves. ^app^V_max_ values are expressed as pmoles/s/mL/µg protein while ^app^K_0.5_ and ^app^EC_50_ as mM. Data are expressed as mean ± standard error of the mean (SEM) of at least three different experiments.

| Conditions | <sup>app</sup>V<sub>max</sub>. | <sup>app</sup>K<sub>m</sub> | <sup>app</sup>K<sub>0,5</sub> | Hill (<sup>app</sup>n<sub>H</sub>) |
| --- | --- | --- | --- | --- |
| Control | 45.07 ± 1.72 | 35.53 ± 4.28 | | |
| 3 mM ADP | 64,19 ± 2,82 | 40.69 ± 5.36 | | |
| 3 mM ATP +<br>regenerating system | 11.29 ± 0.57 | | 59.80 ± 3.51 | 2.24 ± 0.23 |

**Figure 5.**
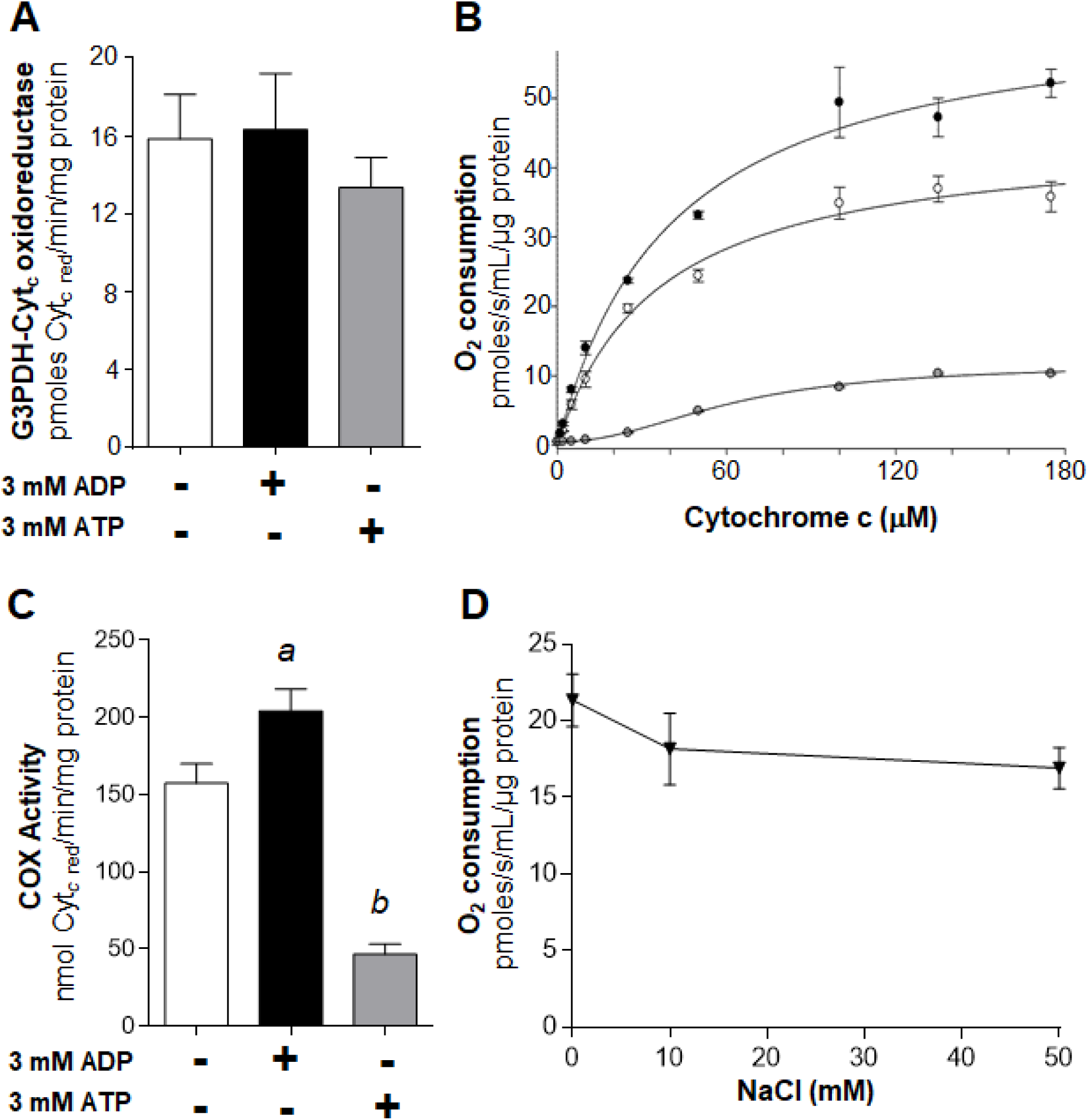
Adenylates specifically and allosterically regulate COX activity in *A. aegypti* flight muscle. (A) G3P-cytochrome *c* oxidoreductase activity was determined in flight muscle homogenates incubated for 30 min at room temperature in the absence (white bar) or in the presence of 3 mM ADP (black bar), or 3 mM ATP (grey bar). G3P-induced reduction of cyt*c*, sensitive to antimycin a was followed spectrophotometrically at 550 nm. (B) Saturation kinetics curves for COX were determined by high resolution respirometry of permeabilized flight muscle using cyt*c* concentrations from 0 to 175 µM in the presence of TMPD-ascorbate plus 3 mM ADP (black circles) or 3 mM ATP + ATP-regenerating system (grey circles) or in the absence of adenylates (white circles). The samples were pre-incubated for 30 min at 27.5°C in the presence of 4 µg/mL oligomycin, 0.5 µM rotenone and 3 mM ADP or 3 mM ATP plus an ATP-regenerating system before the experiments. COX activity was measured in the presence of 2.5 µg/mL antimycin A. (C) COX activity was determined spectrophotometrically in flight muscle homogenate samples incubated for 30 min at room temperature in the absence (white bar) or in the presence of 3 mM ADP (black bar), or 3 mM ATP + ATP-regenerating system (grey bar). Subsequently, 35 µM ferricytochrome *c* was added and COX activity was assessed by following the decrease in absorbance due to the oxidation of cyt*c* at 550 nm. Finally, 1 mM KCN was added to cuvettes to inhibit COX activity, which was considered as the cyanide sensitive rate of cyt*c* oxidation. (D) The effect of ionic strength on COX activity was determined by high resolution respirometry of permeabilized flight muscle using 50 µM cyt*c* in the presence of TMPD-ascorbate and in the absence of adenylate. The samples were pre-incubated for 30 min at 27.5 °C in the presence of 4 µg/mL oligomycin, and NaCl concentrations from 0 to 50 mM. COX activity was measured in the presence of 2.5 µg/mL antimycin A using 2 mM ascorbate and 0.5 mM TMPD as an electron-donor regenerating system. To distinguish cellular respiration from TMPD chemical auto-oxidation, 5 mM KCN was added at the end of each experiment, and COX activity was considered as the cyanide-sensitive rate of oxygen consumption. Comparisons between groups were done by one-way ANOVA and *a posteriori* Tukey’s multiple comparisons tests with ^a^ *p*<0.01 relative to control and ATP, ^b^ *p*<0.05 relative to control. Data are expressed as mean ± standard error of the mean (SEM) of at least three different experiments.

### High energy phosphate recycling mechanisms do not contribute to the regulatory effects of adenylates on COX activity and mitochondrial G3P oxidation

In order to maintain energy homeostasis during active workload, highly oxidative muscles are equipped with mechanisms to allow efficient high energy phosphate transfer from mitochondria to myofibrils [52]. The creatine kinase (CK) shuttle represent such an energy transfer mechanism in mammalian skeletal muscle, while in invertebrates the arginine kinase (ArgK) shuttle has been reported to play similar energetic functions [53,54]. Beyond these mechanisms, adenylate kinase (AK) is also present in mammalian skeletal muscle, catalyzing the reversible ATP synthesis/hydrolysis from/to ADP depending on the cellular adenylate balance [55]. In order to verify the possibility that the regulatory effects of ADP on COX activity would involve a flight muscle AK-mediated conversion of ADP in AMP + ATP, we performed experiments in the presence of 10 µM Ap5a, a classical AK inhibitor [55]. Figure 6A shows that Ap5a caused no apparent effects on COX activity, ruling out the potential contribution of AMP derived from AK as part of the regulatory mechanism observed in *A. aegypti* flight muscle.

**Figure 6.**
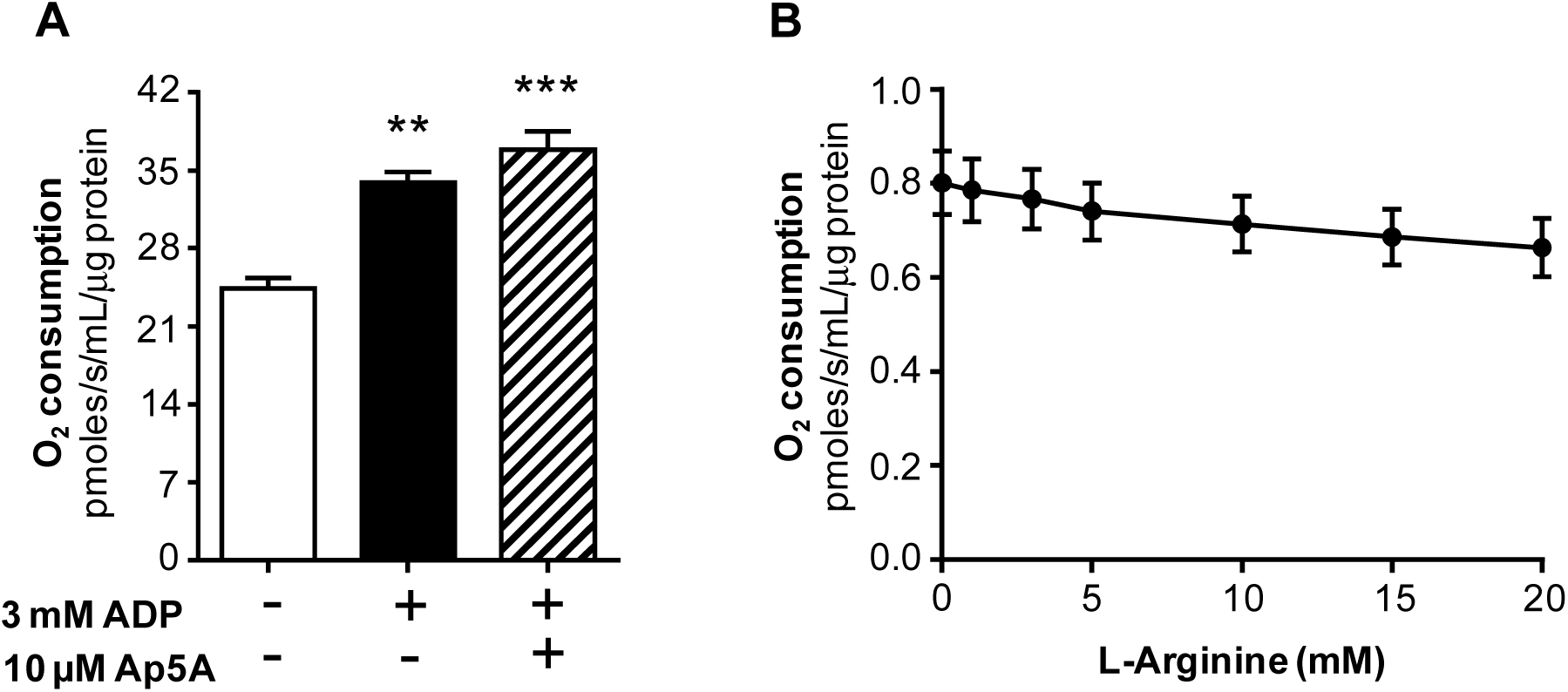
High energy phosphate recycling mechanisms do not contribute to the regulatory effects of adenylates on COX activity or respiration. (A) Assessment of COX activity in mechanically permeabilized flight muscle was carried out in respiration buffer (120 mM KCl, 5 mM KH_2_PO_4_, 3 mM Hepes, 1 mM EGTA, 1.5 mM MgCl_2_, and 0.2 % fatty acid free bovine serum albumin, pH 7.2) containing 1 % Tween 20 at 27.5°C. in the absence (white bars), or in the presence of 3 mM ADP (black bar) or 3 mM ADP + 10 mM Ap5A (hatched bar) to inhibit adenylate kinase. Aliquots corresponding to 4 µg/mL oligomycin, 0.5 µM rotenone, 3 mM ADP, 10 µM Ap5A were added in the respirometer chamber and kept under stirring for 30 minutes. Control identical experiments without Ap5A addition were also performed. Then, 50 µM cyt*c*, 2 mM ascorbate, 0.5 mM TMPD and 5 mM KCN were sequentially added to determine COX activity. (B) The effect of ArgK on flight muscle respiration was evaluated by placing a single insect thorax into the O2K chamber filled with 2 mL of respiration buffer in the presence of 0.5 µM rotenone and 20 mM G3P followed by 3 mM ATP, plus an ATP-regenerating system as described in the methods section. Subsequently, the effect of arginine on respiration was evaluated by stepwise additions from 0 to 20 mM of L-arginine in the respirometer chamber. Data are expressed as mean ± standard error of the mean (SEM) of at least three different experiments.

High energy phosphate transfer mediated by ArgK was previously shown to be functional in insects [54], and its potential regulatory effect on G3P-linked respiratory rates was investigated in *A. aegypti* flight muscle. Figure 6B shows that arginine supplementation to permeabilized flight muscle in the presence of 3 mM ATP and an ATP regenerating system caused any effect on respiratory rates. This strongly indicates that, if present, ArgK activity had a negligible contribution on ADP generation, excluding its involvement on adenylate balance in *A. aegypti* flight muscle.

## Discussion

We described here a regulatory mechanism of mitochondrial G3P oxidation mediated by adenylates which occurs essentially through the allosteric modulation of COX activity, and apparently not at any other component of the electron transport system in the flight muscle of *Aedes aegypti* mosquitoes. To our knowledge, this is the first description of the modulatory effects of mitochondrial G3P oxidation by adenylates and underscores the importance of allosteric regulation of COX activity in a tissue with very high ATP turnover rates. Therefore, beyond the hormonal and ionic mediated effects on G3P oxidation [56], adenylates also emerge as direct regulators of mitochondrial G3P metabolism by allosterically modulating COX activity. Conceivably, allosteric regulation of COX activity by adenylates would play a central role not only to flight muscle energy metabolism, but also to insect dispersal and reproduction. A summary of the regulatory effects of adenylates on mitochondrial G3P oxidation and COX activity in *A. aegypti* flight muscle are outlined in Figure 7.

**Figure 7.**
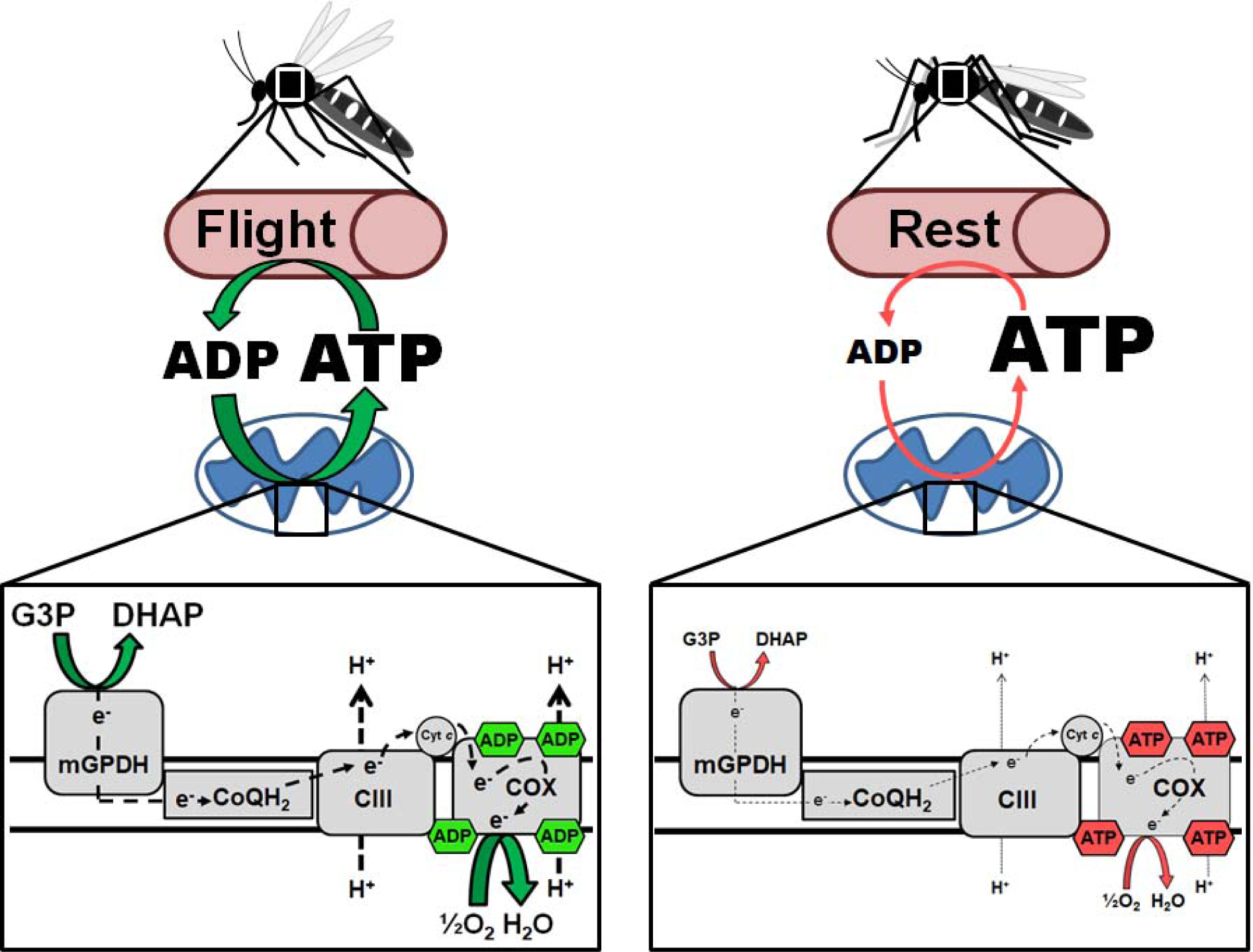
A summary of the regulatory effects of adenylates on COX activity and the impacts on mitochondrial G3P oxidation in *A. aegypti* flight muscle. The flight activity induces a strong increase in energy demand which is reflected by an increase in ADP levels, stimulating COX activity and mitochondrial G3P oxidation. Conversely, when the energy demand posed by flight activity ceases, ADP levels drops and ATP rises, rapidly returning to its pre-flight levels directly impacting COX activity and ultimately affecting mitochondrial G3P oxidation.

Insect flight muscles are highly aerobic tissues, and most of the ATP is made available through a very efficient oxidative phosphorylation process within mitochondria, while most of the demand is posed by key mechanisms for muscle contraction including actomyosin ATPase, Ca^2+^-ATPase and Na^+^/K^+^-ATPase [57]. Thus, insect flight muscles are suitable systems to study the mechanisms underlying respiratory control. Indeed, the extremely high ATP turnover rates found in insect flight muscle can be explained by the huge energy demand posed by flight to allow muscle contraction, causing 50 to 100-fold increases on respiratory rates in the rest - flight transition [5,6]. This large increase in respiratory rates upon initiation of flight implies that *in vivo* there is a tight control of respiration which is not limited by oxygen availability, since flight muscle is enriched with tracheae that deliver molecular oxygen close to mitochondria. In this sense, a structural hallmark of insect flight muscle is the presence of mitochondria with highly packed cristae which establish close interaction with myofibrils [10,58]. In fact, the close interaction between these cell structures strongly suggests a bioenergetic link between the energy supply (mitochondria), and the energy demand sites (myofibrils). Given that AK and ArgK activities seem to play a negligible role on adenylates re-cycling in *A. aegypti* flight muscle (Figure 6), it is plausible that regulation of respiratory rates would be directly mediated by the energy balance between the mitochondrial ATP supply, and the myofibrils ATP demand. Therefore, this mechanism will not require an intermediate process to handle and transfer high energy phosphates between mitochondria and myofibrils, but rather would be exerted directly through adenylates levels and diffusion.

Classically, one of the processes that regulate oxidative phosphorylation is the cellular energy demand, which is mediated by the dissipation of *pmf* by F_1_F_o_ ATP synthase during catalysis [15–17], but also through direct regulation of mitochondrial enzymes [18,21–23,48]. For instance, in different insect species, ADP activates isocitrate and proline dehydrogenases [18,21], by increasing the affinities for their substrates. When *A. aegypti* flight muscle oxidize G3P, ADP increased both maximal uncoupled respiratory rates and the apparent affinity for ADP (Figures 2B, 3, S1, 4A, Tables 1 and 2), while ATP potently inhibited respiratory rates compared to controls (Figures 2B, 4B and Table 2). The activating effects of ADP involve a slightly positive cooperative mechanism and are independent of the substrates used (Figures 2 and S1). Considering that maximal respiratory rates were measured in the presence of a proton ionophore (FCCP), we concluded that regulatory effects of adenylates act independent of the *pmf*. However, the magnitude of respiratory activation by ADP varied largely on the fuel preference (Figures 2 and S1). We interpreted this differential effect by the ability of ADP to boost the activity of a number of mitochondrial dehydrogenases which are necessary to allow respiration linked to pyruvate and proline metabolism, but not linked to G3P oxidation [18,21]. Indeed, using complex I-linked substrates, respiratory rates increased ∼ 10 times, while G3P-linked respiration increased by ∼ 45% upon ADP exposure (Figures 2 and S1). This indicates that, beyond their known activating effects on mitochondrial dehydrogenases, ADP also activates respiration through a mechanism independent of substrate transport across the mitochondrial inner membrane, TCA cycle, and *pmf*. Two possibilities arise as the potential mechanisms by which adenylates regulate respiratory rates: the mG3PDH–cytochrome *c* (mG3PDH-cyt*c*) oxidoreductase, and COX activities. Adenylates were ineffective in regulating mG3PDH-cyt*c* oxidoreductase activity (Figure 5A), strongly suggesting that none of the electron transport system components from mG3PDH to cyt*c* are targeted by adenylates. Indeed, adenylates caused strong regulatory effects on COX activity in *A. aegypti* flight muscle (Figures 5B, 5C and Table 3). COX activity can be regulated by different factors, including reversible phosphorylation [59–61], interaction with other proteins, assembly as supercomplexes [62], sodium and calcium ions [63] and by adenylates [22,23,32–35]. From the 10 sites of ADP binding to COX identified in bovine heart, seven are exchanged by ATP in conditions of low energy demand [22,32,33]. While subunit IV binds ADP/ATP at both sides of mitochondrial inner membrane, subunit VIa binds adenylates only at the matrix side [23,35]. Indeed, the experiments shown here demonstrate that the activating effects of ADP on respiration are partially reversed by CAT, indicating the involvement of two distinct components that mediate this regulatory effect: one in the matrix, and the other at the intermembrane space (Figure 4A and Table 2). Despite the use of TMPD-ascorbate system was reported to inhibit the cooperativity of the two cyt*c* binding sites in the dimeric COX complex, preventing the binding and dissociation of cyt*c* in the presence of ATP in mammalian models [22,64], sigmoidal COX kinetic curves were maintained by increasing ATP levels in *A. aegypti* flight muscle (Figure 5B). In this regard, changes in COX kinetics from hyperbolic to sigmoidal shaped curves induced by ATP, can result from cooperative interaction between the two cyt*c* binding sites in the monomeric COX, or eventually by interaction between COX dimers [22,23,25,65]. Potentially, given that in our experiments we used mechanically permeabilized tissue that conserves mitochondrial morphology in the cellular environment, and preserve their contacts with other organelles, COX kinetics might slightly differ from the isolated enzyme complex.

During the rest-flight transition, the huge increase in energy demand in the flight muscle stimulates respiration and decreases the ATP/ADP ratio. Although quantification of adenylates content is difficult due to rapid ATP hydrolysis during extraction, total adenylate levels around 6 mM were estimated in mammalian heart and skeletal muscle [66] and in blowfly flight muscle [18,67], which are within the range of ADP and ATP used in our experiments here. However, even during forced flight activity, ATP represents the dominant form of adenylates in blowflies [68], suggesting that other mechanisms beyond direct control of enzyme activity should take place *in vivo*. Potential explanations for this apparent paradox, might involve the different levels of adenylates in cellular compartments, as in mitochondrial matrix ATP/ADP ratios are ∼ 10, while in cytosol is ∼1000 [69]. Therefore, maintenance of such lower ATP/ADP ratios in mitochondria would favor higher respiratory rates through the activation of mitochondrial enzymes, including COX. In addition, the rapid diffusion of adenylates through the cell might also play an important role in regulating respiratory rates, especially in the case of insect flight muscle where mitochondria establish close interactions with myofibrils [10] potentially facilitating the exchange of ATP and ADP bypassing their simple diffusion through the cytosol. Finally, post-translational modifications of mitochondrial enzymes, including cAMP-dependent protein kinase (PKA)-mediated phosphorylation of COX [59–61], modulate its activity according to changes in cellular energy balance. Indeed, phosphorylation of specific COX subunits at serine residues might either potentiate (subunits II, III and Vb [59], or prevent the inhibitory effects of ATP (subunit IV-1 [60]). Conceivably, when cellular ATP consumption is high, substrate oxidation through the TCA cycle produces high levels of CO_2_, which in turn stimulates COX activity by preventing the inhibitory effects of ATP [61]. Conversely, when the energy demand posed by flight activity ceases, the ATP content rapidly return to its pre-flight levels, owed the extreme capacity of flight muscle oxidative phosphorylation, directly affecting respiratory rates by inhibiting COX. The powerful inhibitory effect of ATP observed in *A. aegypti* flight muscle (Figures 2B, 4B, 5B and 5C) can be explained by one of the following mechanisms: *i)* the close interactions between myofibrils and mitochondria, which allow rapid ATP accumulation near and inside mitochondria, bypassing diffusion; *ii)* the absence of high energy phosphate shuttle such as ArgK and AK in a way that it would make it more susceptible to the inhibitory effects of ATP (Figures 4B, 5B, and 5C); *iii)* different amino acid composition of the allosteric ATP binding site in COX subunit IV in insects making it less sensitive to phosphorylation induced by the PKA. Finally, one might consider that regulation of respiratory rates in insect flight muscle should be tighter than in mammalian heart for example, as insect flight is not a continuous biological task, which is not the case of the beating heart, and thus can be strongly down-regulated according to the biological needs for dispersal. Given the huge capacity of flight muscle oxidative phosphorylation to produce ATP, strong inhibition of COX activity by ATP may represent an ingenious strategy developed by insects to spare the available nutrients in rest conditions for use only when energy demand of flight is posed. In this sense, this allosteric inhibitory effect of ATP resembles that existent in phosphofructokinase I and pyruvate kinase, where ATP exerts strong inhibitory control on glycolytic flux, thus avoiding unnecessary nutrient utilization [70,71].

Previous evidence from our laboratory revealed that respiration in *A. aegypti* flight muscle is essentially sustained by means of pyruvate, proline and G3P oxidation [11]. G3P metabolism involves the action of specific cytosolic and mitochondrial dehydrogenases (cG3PDH and mG3PDH), which compose the glycerophosphate shuttle that transfer reduced equivalents from the cytosol to electron transport system [56]. The glycerophosphate shuttle represents a key intersection between glycolysis, lipogenesis and oxidative phosphorylation [56], and specific regulation of G3P oxidation by adenylates might have important consequences to flight muscle metabolic homeostasis. Conceivably, in conditions where ATP levels rise, such as in resting flight muscle, G3P will be spared in this tissue and can be diverted out of flight muscle to allow phospholipid and triglyceride biosynthesis in the fat body and ovaries for energy storage and reproduction purposes. Also, since insect flight muscle has a very active glycolytic pathway, it is possible that activation of glycerophosphate shuttle would play a key metabolic role by providing a continuous source of NAD+ to sustain high glycolytic flux [56], while is spared of intense acidification generated by lactate dehydrogenase for the same purpose [40,41]. Interestingly, the activities of enzymes involved in the glycerophosphate shuttle remarkably correlated with the glycolytic capacity of insect flight muscles [72], suggesting a direct connection between G3P and glucose metabolism. Potentially, modulation of mitochondrial G3P oxidation by adenylates through allosteric regulation of COX, may represent a novel way to control glucose and lipid metabolism in different insect tissues that ultimately affect reproduction and dispersal.

In conclusion, we demonstrate here that mitochondrial G3P oxidation in *A. aegypti* flight muscle is regulated by adenylates through allosteric modulation of COX activity. The observations reported here represent a beautiful example of co-adaptation between the energy supplying and demanding processes through the modulation of COX activity by adenylates, expanding our view on the regulation of oxidative phosphorylation in a system with very high metabolic rates, and with potential consequences to mosquito metabolism and biology.

## Materials and methods

### Insects

*Aedes aegypti* (Red eyes strain) were maintained at 28 °C, 70–80 % relative humidity with a photoperiod of 12 h light/dark (L:D, 12:12 h). Larvae were reared on a diet consisting of commercial dog chow. Adults were kept at the same temperature, humidity, and photoperiod. Insects utilized in all experiments were female individuals, 5-7 days after the emergence. Usually about 200 insects were placed in 5 L plastic cages in a 1:1 sex-ratio and fed *ad libitum* on cotton pads soaked with 10 % (w/v) sucrose solution.

### Isolation of mitochondria

Mitochondria isolation from *A. aegypti* flight muscle was performed as previously described by our group [11]. Protein concentration was determined by the Lowry method, using bovine serum albumin as standard [73].

### Respirometry analyses

The respiratory activity of mechanically permeabilized flight muscle and isolated mitochondria from *A. aegypti* was carried out in a two-channel titration injection respirometer (Oxygraph-2k, Oroboros Instruments, Innsbruck, Austria) at 27.5 °C using methods previously established by our group [11,74]. Two distinct substrate combinations were used in our experiments as following: pyruvate + proline (Pyr+Pro) or *sn*-glycerol 3-phosphate (G3P) in presence and absence of ADP. The routine was started by addition of the substrates to final concentrations of 10 mM Pyr + 10 mM Pro or 20 mM G3P. When using G3P, 0.5 µM rotenone was added before substrate addition to avoid electron backflow from G3P dehydrogenase (G3PDH) to complex I. The ATP synthesis coupled to oxygen consumption was induced by 1 mM ADP followed by a second shot reaching a final ADP concentration of 2 mM. Then, 10 μM cyt*c* were added to each respirometer chamber and the absence of a significant stimulatory effect on respiration was used as a quality control test for integrity of the outer mitochondrial membrane [50]. Control experiments in the absence of ADP were also performed. Maximum uncoupled respiration was induced by stepwise titration of carbonyl cyanide *p*-(trifluoromethoxy) phenylhydrazone (FCCP) to reach final concentrations of 2.0 µM or 2.5 µM for isolated mitochondria and whole flight muscle, respectively. Finally, respiratory rates were inhibited by the addition of 2.5 µg/mL antimycin A. The maximal respiratory rates (ETS) from each experiment were calculated by subtracting the antimycin resistant oxygen consumption from FCCP-stimulated oxygen consumption rates. ADP saturation kinetic curves were determined polarographically in mechanically permeabilized flight muscle and in isolated mitochondria. The experiment was started by the addition of 0.5 µM rotenone and 20 mM G3P. After stabilization of oxygen consumption, 10 μM cyt*c* followed by 2.0-2.5 µM FCCP were added, and then ADP titration was carried out by stepwise additions from 0 to up to 10 mM to induce maximal respiratory rates. Finally, respiratory rates were inhibited by adding 2.5 µg/mL antimycin A. The contribution of the adenine nucleotide translocator (ANT) activity on ADP regulatory effects on respiration was determined by adding 50 µM carboxyatractyloside (CAT) before ADP titration. Regulation of respiratory rates by ATP was assessed in mechanically permeabilized flight muscle in the absence of ADP, by performing stepwise titration of ATP from 0 to 10 mM in the presence of an ATP-regenerating system consisting of 5 mM phosphenolpyruvate, 5 mM MgSO_4_, 20 U/mL pyruvate kinase, 30 U/mL lactate dehydrogenase and 0.2 mM NADH [75]. In order to evaluate the role of the arginine kinase (ArgK) in the regulation of flight muscle respiration, a single insect thorax was added into the O2K chamber filled with 2 mL of respiration buffer. Then, 0.5 µM rotenone and 20 mM G3P followed by 3 mM ATP, plus an ATP-regenerating system described above, were added. The contribution of ArgK on respiration of mechanically permeabilized flight muscle was evaluated by stepwise additions from 0 to 20 mM of L-arginine in the respirometer chamber. To evaluate the effect of ionic strength in the regulation of flight muscle respiration, a single insect thorax was added into the O2K chamber filled with 2 mL of respiration buffer. Then, 0.5 µM rotenone, 20 mM G3P and 2.5 µM FCCP were added. The effect of ionic strength on respiration of mechanically permeabilized flight muscle was evaluated by stepwise additions from 0 to 50 mM of NaCl in the respirometer chamber. To assess respiratory capacity in mechanically permeabilized flight muscle, all experiments were performed in an environment enriched with oxygen within the O2K chamber, as recommended by the literature [50,76]. Experiments started by injecting a suitable amount of oxygen-enriched gaseous mixture (70% O_2_ and 30 % N_2_ mol/mol) into the O2K chamber to reach an oxygen concentration close to 500 nmol/mL. To avoid significant oxygen depletion, and the potential side effects on respiratory rates, oxygen injections were performed once the oxygen concentration fell down below 150 nmol/mL into the O2K chamber.

### Flight muscle homogenate

Five thoraces from adult females were homogenized in a ground glass potter with 500 µL of hypotonic buffer (25 mM potassium phosphate and 5 mM MgCl_2_, pH 7.2). Subsequently the solution was centrifuged at 1500 x *g* for 10 minutes, the supernatant was recovered, and the protein content was determined by the Lowry method [73].

### Mitochondrial G3P-cyt*c* oxidoreductase activity

The mitochondrial G3P dehydrogenase-cytochrome *c* (mG3PDH-cyt*c*) oxidoreductase activity was determined by assessing the increase in absorbance at 550 nm due to the reduction of ferricytochrome *c* induced by the oxidation of G3P at room temperature, in 1 mL of respiration buffer, pH 7.2 using a Shimadzu spectrophotometer model 2450 (Shimadzu Scientific Instruments, Tokyo, Japan). The experiments were conducted using samples corresponding to 70 μg of proteins from six different insect cohorts. To assess the effect of adenylates, the samples were incubated for 30 min at room temperature in the presence of 1 mM KCN, 0.5 μM rotenone and 3 mM ADP or 3 mM ATP plus the ATP-regenerating system, before measuring the enzymatic activity. Afterwards, the reaction was initiated by the addition of 50 μM ferricytochrome *c* followed by 20 mM G3P and the absorbance was monitored at 550 nm for about 5 minutes. Antimycin A (2.5 μg/mL) was added to inhibit complex III activity and the mG3PDH-cyt*c* oxidoreductase activity was considered as the antimycin A-sensitive rate of cyt*c* reduction (ε = 18.7 mM ^1^ • cm ^1^). Control experiments in the absence of adenylates were also performed.

### COX activity

COX activity was measured polarographically using the high-resolution Oxygraph-2k system (Oroboros, Innsbruck, Austria). A single *A. aegypti* mechanically permeabilized flight muscle from a female was transferred into the O2K chamber containing 2 mL of the respiration buffer with 1 % Tween 20 (v/v). Then the sample was incubated for 30 min at 27.5 °C in the presence of 4 µg/mL oligomycin, 0.5 µM rotenone and 3 mM ADP or 3 mM ATP plus the ATP-regenerating system described above. Control experiments in the absence of adenylates were also performed. COX activity was measured in the presence of 2.5 µg/mL antimycin A and various concentrations of cyt*c* (0-175 μM) using 2 mM ascorbate and 0.5 mM N,N,N’,N’-Tetramethyl-*p*-phenylenediaminedihydrochloride (TMPD), as an electron-donor regenerating system. To distinguish cellular respiration from TMPD chemical auto-oxidation, 5 mM KCN was added at the end of each experiment, and COX activity was considered as the cyanide-sensitive rate of oxygen consumption. To verify the possibility that COX activation by ADP would involve adenylate kinase (AK) mediated conversion of ADP in AMP + ATP, experiments in the presence of Ap5a, a classical AK inhibitor [55] were performed. The samples were incubated for 30 minutes on the respirometer chamber filled with 2 mL of respiration buffer with 1 % Tween 20 (v/v) in the presence of 4 µg/mL oligomycin, 0.5 µM rotenone, 3 mM ADP and 10 µM Ap5A. Then, 50 µM cyt*c*, 2 mM ascorbate, 0.5 mM TMPD and 5 mM KCN were sequentially added to determine COX activity. Control identical experiments without Ap5A addition were also performed. Since COX activity is limited by low oxygen concentrations [76,77], the oxygen enriched gas mixture (70% O_2_ and 30 % N_2_ mol/mol) was also injected to reach an oxygen concentration of about 500 nmol/mL before measuring enzyme activity.

Alternatively, COX activity was determined spectrophotometrically in flight muscle homogenate samples using a Shimadzu spectrophotometer model UV-2450 (Shimadzu Scientific Instruments, Tokyo, Japan) following methods described in the literature with slight modifications [78]. The experiments were conducted at room temperature, in 1 mL of respiration buffer using samples corresponding to 10 μg of proteins from four different insect cohorts. To assess the effect of adenylates, the samples were incubated for 30 min at room temperature in the presence of 2.5 μg/mL antimycin A and 3 mM ADP or 3 mM ATP plus the ATP-regenerating system, before measuring the enzymatic activity. Afterwards, the reaction was initiated by the addition of 35 μM reduced ferricytochrome *c* and the absorbance was monitored at 550 nm for about 5 minutes. Afterwards, a volume corresponding to 1 mM KCN final concentration was added to cuvettes to inhibit COX activity, which was considered as the cyanide sensitive rate of cyt*c* oxidation (ε = 18.7 mM ^1^ • cm ^−1^). Control experiments in the absence of adenylates were also performed.

The effect of ionic strength on COX activity was investigated by high resolution respirometry of permeabilized flight muscle using 50 µM cyt*c* in the presence of TMPD-ascorbate and in the absence of adenylates. The samples were pre-incubated for 30 min at 27.5 °C in the presence of 4 µg/mL oligomycin, and NaCl concentrations from 0 to 50 mM. COX activity was measured in the presence of 2.5 µg/mL antimycin A using 2 mM ascorbate and 0.5 mM TMPD as an electron-donor regenerating system. To distinguish cellular respiration from TMPD chemical auto-oxidation, 5 mM KCN was added at the end of each experiment, and COX activity was considered as the cyanide-sensitive rate of oxygen consumption.

### Kinetic and data analyses

To determine the kinetic parameters of respiratory rates and COX activity, fits were performed using the kinetics module of Sigmaplot 11 for Windows Regression Wizard (Systat Software, Inc., USA). For the ADP activation of respiratory rates in all states, after several rounds of fitting with the most known kinetic models, the approach with the non-linear regression dynamic fitting using a Hill equation with four parameters, according to equation 1,

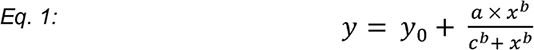

resulted in the best solution, with values of R^2^>0.99 in all cases, where *b* is the apparent Hill coefficient of cooperativity (^app^n_H_), a is apparent V_max_, c is apparent K_0.5_ and x the substrate concentration. For the ATP inhibition of respiratory rates, a non-linear regression using the standard curve, five parameter logistic equation with two slopes (Equation 2), gave the best fit with R^2^>0.99 and for this reason was employed:

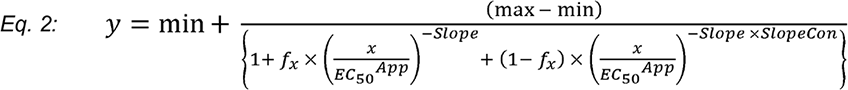

Where 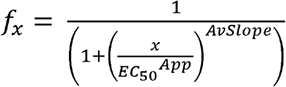 and 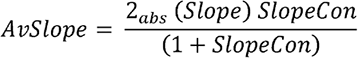

To determine the effect of ADP on COX activity kinetic parameters, the best solution for control and ADP curves were obtained using classical Michaelis–Menten equation (equation 3), with values of R^2^>0.99, while for ATP curves, the Hill equation with four parameters (equation 1), revealed the best fitting option, with R^2^ ∼ 0.998.

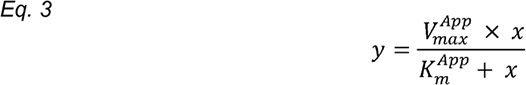

In Michaelis–Menten equation 3 above, x represents the substrate concentration. Apparent kinetic parameters (^app^V_max_, ^app^K_0.5_, ^app^n_H_, ^app^EC_50_, ^app^K_m_) were calculated assuming the best fitting equation described above and were presented as mean ± standard error of the mean (SEM) of at least four different experiments (n≥4). Data in all graphs were presented as mean ± SEM of values for each condition. D’Agostino and Pearson normality tests were done for all experimental groups to assess their Gaussian distribution. Comparisons between groups were done by unpaired Student’s t-test, Mann Whitney’s test, one-way ANOVA and *a posteriori* Tukey’s multiple comparisons test, and differences with *p*<0.05 were considered significant. Graphs were prepared by using the GraphPad Prism software version 6.00 for Windows (GraphPad Software, USA) or Sigmaplot 11 for Windows (Systat Software, Inc., USA).

## Abbreviations

G3P: *sn*-glycerol 3-phosphate
G3PDH: glycerol 3-phosphate dehydrogenase
Pyr: pyruvate
Pro: proline
FCCP: carbonyl cyanide *p*-(trifluoromethoxy) phenylhydrazone
AK: adenylate kinase
ArgK: arginine kinase
CAT: carboxyatractyloside
KCN: potassium cyanide
*pmf*: protonmotive force
TMPD: N,N,N’,N’-Tetramethyl-*p*-phenylenediaminedihydrochloride
COX: cytochrome *c* oxidase
ANT: adenine nucleotide translocator
Ap5a: P1,P5-di(adenosine-5’)pentaphosphate
cyt*c*: cytochrome *c*

## Author contributions

AG, JBRCS, JAM, CFLF and MFO conceived and designed the experiments. AG and JBRCS performed the experiments. JAM and CFLF performed the kinetic analyses. AG, JBRCS, JAM, CFLF and MFO analyzed and interpreted the data. AG, JAM, CFLF and MFO contributed reagents/materials/analysis tools. AG, JAM, CFLF and MFO wrote the paper; all authors had approval of manuscript.

## Acknowledgments

We would like to thank Mrs. Jaciara Miranda Freire for the excellent technical assistance on maintenance of *A. aegypti* colony. This work was supported by grants from Conselho Nacional de Desenvolvimento Cientifico e Tecnológico (CNPq) [#404153/2016-0 MFO, and 483334/2013-8 AG], and Fundação Carlos Chagas Filho de Amparo à Pesquisa do Estado do Rio de Janeiro (FAPERJ) [#E-26/102.333/2013, E-26/203.043/2016, and E-26/111.169/2011]. AG, MFO and CFLF are fellows from CNPq [#402409/2012-4, 303044/2017-9 and 308847/2014-8]. JBRCS and AG were fellows from PAPD-FAPERJ [#E-26/102.752/2011 and E-44/208702/2014].

