## Supplemental Figure 1 for "Mitochondrial glycerol phosphate oxidation is modulated by adenylates through allosteric regulation of cytochrome *c* oxidase activity in mosquito flight muscle"

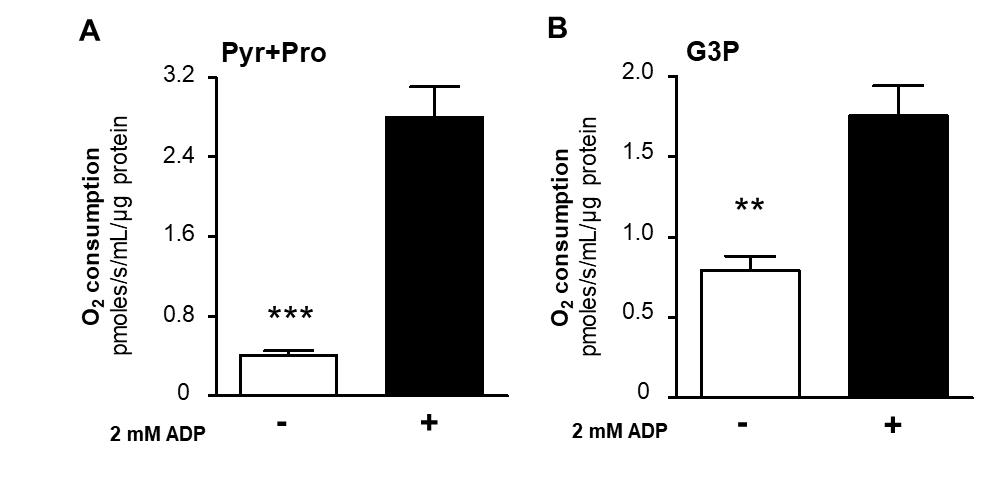

**Figure S1. Adenylates regulate respiratory rates by a mechanism independent of substrate transport to mitochondria, tricarboxylic acid cycle enzymes activities, and protonmotive force.** Maximum uncoupled respiratory rates (2.5 μM FCCP) of isolated mitochondria from adult females were determined by high resolution respirometry in the absence (-) or in the presence (+) of 2 mM ADP using 10 mM pyruvate plus proline (A), or 20 mM G3P (B) as substrates. When using G3P, 0.5 µM rotenone was added before G3P to avoid electron backflow from G3PDH to complex I. Data are expressed as mean O_2_ consumption rates (pmoles/s/mL/μg protein) ± standard error of the mean (SEM) of at least seven different experiments. Comparisons between groups were done by unpaired Student´s t test or Mann Whitney’s test with ****p*<0.0001, ***p*<0.005 relative to control group (without ADP).
